# Performance of a winter wheat composite cross population in two temperate agroforestry systems – a case study

**DOI:** 10.1101/2023.02.07.527524

**Authors:** Christina Vaccaro, Fabio Mascher, Johan Six, Christian Schöb

## Abstract

In agroforestry systems (AFS), where environmental conditions are highly variable at small spatial scales, the use of uniform genetic material of a single cultivar commonly grown in monoculture cropping might not be optimal. However, the use of composite cross populations (CCPs) that contain an inherent genetic variability might be a promising approach to match the environmental variability created by trees in AFS. In this experimental trial, the performance of a CCP (‘CC-2k’) of winter wheat was compared to a commercial variety (‘Wiwa’) in a split-plot design in two AFS (Feusisberg, Wollerau) in Central Switzerland. The factor variety (p < 0.05) and the interactions of site and distance to tree (p < 0.05) and site and variety (p < 0.05) significantly affected wheat yield at the plot level. CC-2k and Wiwa yield (across all distances) amounted to 1.9 ± 0.7 Mg ha^−1^ and 2.0 ± 0.8 Mg ha^−1^, respectively, in Feusisberg and to 1.9 ± 0.9 Mg ha^−1^ and 0.7 ± 0.4 Mg ha^−1^, respectively, in Wollerau. Wiwa had a higher protein, Fe and Ca content than CC-2k. Therefore, while the CC-2k outperformed the commercial organic variety Wiwa in terms of yield in one of the two AFS, Wiwa outperformed CC-2k in terms of quality. The composite cross population might be better adapted to the heterogenous environment of agroforestry systems but fails to reach the high-quality product of modern elite cultivars. Further improvements in terms of quality might make CCPs highly interesting for diversified agricultural systems with larger environmental heterogeneity than common monoculture cropping systems.

## Introduction

In the face of climate change and the ongoing biodiversity crisis – requiring agriculture to adapt to a shifting climate and to be more environmentally-friendly – the development of sustainable and resilient food production systems with adaptive cultivars is highly relevant to meet the Sustainable Development Goals (Bourke *et al*., 2021; Obrecht *et al*., 2021; Rist *et al*., 2020). Agroforestry – agriculture with trees – has gained renewed economic and environmental interest in the past decade as numerous scientists have declared agroforestry a promising approach to regenerative agriculture that conserves and restores soils and biodiversity and potentially offers greater resilience to climate change stressors (e.g., Kohli *et al*., 2007; Jose 2009; Torralba *et al*., 2016). In addition to the inherent interspecific diversity in agroforestry, intraspecific diversity in form of genetic diversity in tree and crop varieties can further contribute to agrobiodiversity and thus to food security (Lin, 2011), health benefits (Fanzo *et al*., 2013), preservation of a rich agricultural knowledge (Morton, 2019) and protection of ecosystem services (Duru *et al*., 2015).

Since the “Green Revolution” in the 1950s, local crop varieties were replaced by standardized, commercial varieties worldwide. For instance, 90% of 10,000 wheat varieties cultivated in China in 1940 were lost in 1970 (Rist *et al*., 2020). How can plant breeders regain intraspecific crop diversity? One approach are composite cross populations (CCPs). Composite crosses are populations obtained by the reciprocal intercrossing of genotypes and mixing of the progeny (Suneson, 1956; Knapp *et al*., 2020). A CCP is therefore a particularly plastic and adaptable culture where each plant constitutes a unique genotype.

A new perspective for the development of agroforestry is the use of CCPs. Their dynamic gene pool provides the potential of adaption to locally heterogeneous environmental conditions found in agroforestry systems (AFS) (Smith, Pearce and Wolfe, 2012a; Bourke *et al*., 2021). Genotypes which are better adapted to local conditions should have more progeny and thus, over time, increase in frequency (Döring *et al*., 2011). The spatial heterogeneity in AFS is created by various tree-crop-interactions dependent on the proximity to a single tree or the tree row, respectively, and the overall-design of the AFS. Aboveground interactions include microclimatic modification (shade, decreased temperature and wind speed, increased water vapor content) and increased insect density and diversity; belowground interactions include competition for water and nutrients or nutrient capture and sharing, hydraulic lift and allelopathy (Jose, Gillespie, and Pallardy 2004). Optimal crop growth in the understorey of an AFS might theoretically require different cultivars that vary in light, water and nutrient requirements. Precision farming at such a small scale is practically challenging. Previous studies suggest that CCPs can provide similar yields and quality as pure varieties with increased yield stability (Döring *et al*., 2015; Vollenweider *et al*., 2020).

The cultivation of CCPs in agroforestry could not only enhance agrobiodiversity, food security, and resilience, but also buffer varying crop yields in AFS. This is desirable because although there is good scientific evidence that AFS increase land use efficiency in many cases (e.g., Dupraz *et al*., 2018a; Graves *et al*., 2007), harvests of the individual crops in AFS tend to show lower yields (in particular close to the tree rows) in temperate climate (Dufour *et al*., 2013; Pardon *et al*., 2018; Carrier *et al*., 2019; Swieter *et al*., 2019).

The use of CCPs in AFS, as suggested by Smith, Pearce, and Wolfe (2012b) seems a promising approach to better match the heterogenous and varying environmental conditions in AFS than varieties bred for high-input and mono-cropping agriculture. Breeding and effective selection of alleles connected to maximum and/or stable yield, respectively, in competitive, heterogeneous and low-input environments as often found in AFS should ideally be undertaken under exactly such conditions. The best varieties specifically targeted for those environments are likely to be produced in breeding programs undertaken in such conditions (Atlin and Frey, 1989).

Given the heterogeneous environment that the implementation of trees create, we hypothesize that a wheat CCP is more apt to cope with heterogeneous environmental conditions in the understorey of trees in a temperate agroforestry system and will therefore outperform a commercial cultivar monoculture in terms of yield and quality in particular in proximity to the tree row.

## Materials and methods

### Experimental Sites

Field experiments were carried out between autumn 2021 and summer 2022 at two organically managed AFS in Feusisberg and Wollerau in Switzerland. Both farms are located in the Kanton Schwyz in Central Switzerland, lying on the upper Lake Zurich. A silvoarable AFS with standard fruit trees (apple, pear, plum) and walnut trees arranged in west-east orientated tree rows is located in Wollerau (N 47°10’53.928’’, E 8°43’56.188’’) on a farm with 4.5 ha. The crop rotation at the experimental field includes lupin, barley, oat, wheat triticale with clover undersown and wheat. The soil is a medium-heavy, slightly acidic loam. The experimental field lies 620 m above sea level on the Northwest side of 16 years old standard apple trees. Standard plum trees are also arranged in west-east orientated tree rows in a natural meadow in Feusisberg (N 47°10’53.928’’, E 8°43’56.188’’). The whole farm size amounts to 14 ha. The soil is a sandy, slightly alkaline loam rich in humus. The experimental field lies 705 m above sea level on the North side of a 30-year-old plum tree row. Both farms’ climate are classified as warm and temperate (“Cfb” in Köppen and Geiger classification) with a mean annual temperature of 8.1°C and a mean annual precipitation of 1618 mm with high amounts of precipitation even in the driest month (February, 12 rainy days) (https://de.climate-data.org, 2022). May has the highest average number of rainy days per month (18 days). Soil carbon, nitrogen and total phosphorous content amounted to 2.99% ± 0.50, 0.34% ± 0.04 and 1000 ± 104 mg P kg^-1^ for Wollerau and 5.83% ± 1.06, 0.58% ± 0.09 and 935 ± 129 mg P kg^-1^ for Feusisberg, respectively, and corresponded to C, N and P levels averagely present in agricultural soils in Switzerland (C: 3.13%, N: 0.29%, P: 932 mg kg^-1^, source: NABO, personal communication), with the exception of higher C and N levels in Feusisberg. The C:N ratio for Wollerau was 8.8, for Feusisberg: 10.2.

### Plant Material

A common winter wheat variety (‘Wiwa’) and a winter wheat CCP (‘CC-2k’) were used in the field experiment: The variety Wiwa is an obtention of the plant breeder Getreidezüchtung Peter Kunz (gzpk) in Hombrechtikon, Switzerland. Released in 2005, Wiwa has been ranked to Swiss baking class TOP (organic class 1) because of its elevated baking properties (Dierauer, 2020) and is recommended for cultivation in organic agriculture. CC-2k is a composite cross obtained as a diallel cross of 20 wheat varieties and breeding lines from Switzerland and from Europe (A. Schori and V. Michel, personal communication). The composite cross population has been constituted in the year 2000 and has been cultivated on 10×10 m plots (100 m^2^) surrounded with a 5 m triticale buffer, during 8 years to obtain the F10 generation (F1012600900), at the Agroscope site in Changins (Nyon, Switzerland). For the present experiments, seed material has been produced with seeds from generation 8 on a 150 m^2^ plot to obtain generation 11 (F1112600900).

### Experimental Design

The experiments were conducted at two trial sites in Feusisberg and Wollerau, respectively. Wiwa and CC-2k were planted in two 1.8 × 9 m strips (“long plot”) in a perpendicular manner on the North side of a tree row. The Wiwa and CC-2k long plots were planted at 2.5 m from the trunk of a reference tree in a “split-plot” design with three replicates per site. Within each long plot, four observation and sampling plots of 1 m^2^ were defined at 1, 2.4, 3.8 and 7 m distance from the tree-row border (Fig. 1). Photosynthetically active radiation (PAR) sensors (Apogee QSO-S PAR Photon Flux, METER Environment) were installed on 27 February 2022 in three replicates (one in Feusisberg, two in Wollerau) at three positions per replicate, i.e. three distinct distances from the tree row. Sensors were placed at 1.3 m height (see SI III) and connected to ECHO Em50 Dataloggers. PAR was measured every 10 minutes as Photosynthetically Active Photon Flux Density (PPFD) in μmol m^-2^ s^-1^ until harvest.

**Fig. 1:**
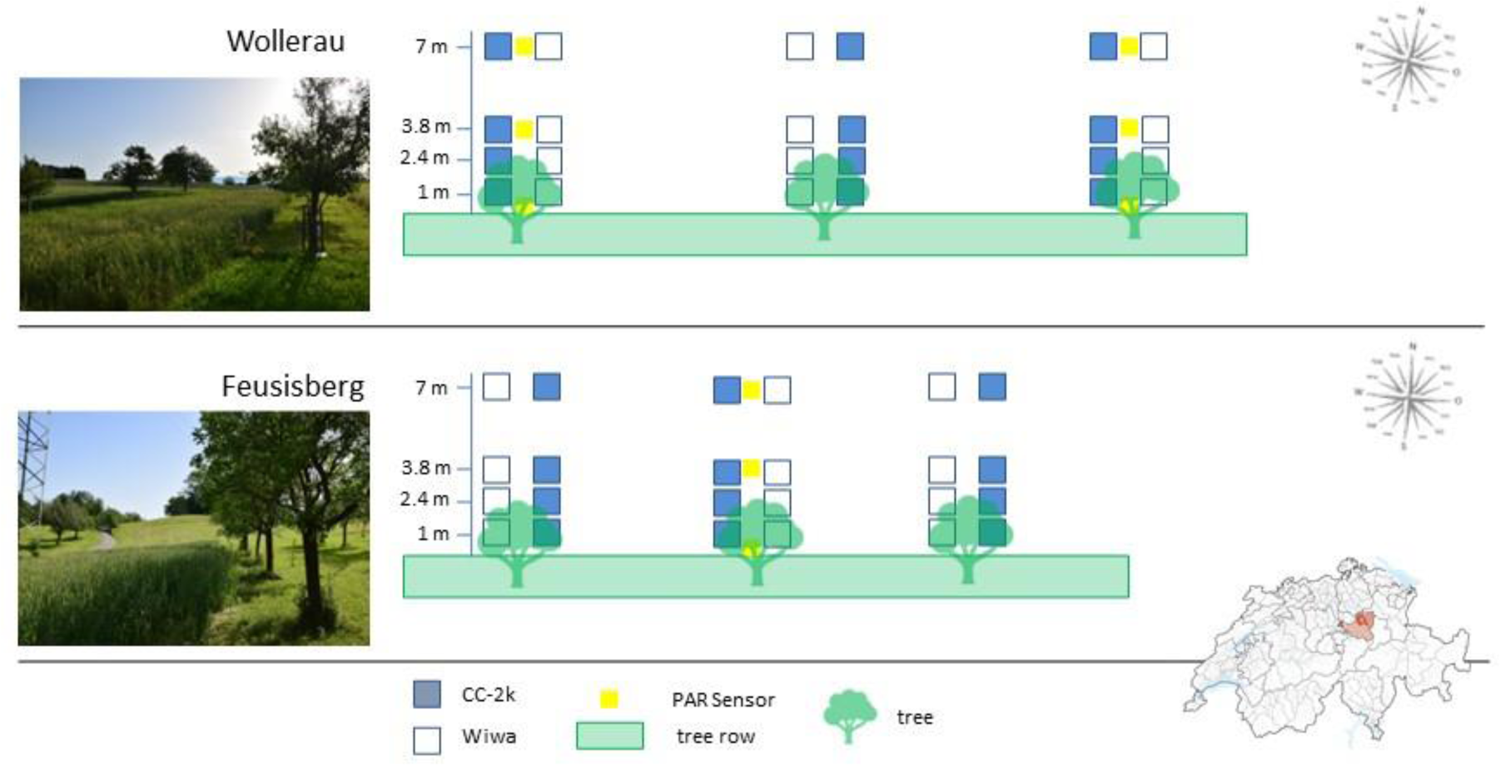
Experimental design at two Swiss agroforestry systems in Feusisberg and Wollerau. Each replicate consisted of a strip (“long plot”) with the composite cross population ‘CC-2k’ and commercial variety ‘Wiwa’. Physiological and yield parameters were collected from 1 m^2^ plots at four distances (1, 2.4, 3.8 and 7 m) from the tree row.

Soil preparation & sowing: The field was tilled with a plough and a harrow in the first week of October 2021. Wiwa and CC-2k were both sown on 15 October 2021 with 350 grains per m^2^ using a small drag coulter seed drill, available at the farm.

Fertilisation was carried out according to farmer’s practice with compost (SI Tab. 2). At the Wollerau site, additional fertilisation was necessary as plants showed yellowing and chlorophyll measurements showed a deficient nutrient status. On 5 May 2022 about 400 hl ha^-1^ of 1:2 diluted liquid manure (cattle slurry with low fecal matter and urine manure diluted) was spread evenly by hand with a hose on the experimental area.

### Plant Measurements and Sampling

Leaf chlorophyll content was assessed indirectly by usage of a chlorophyll meter (SPAD-502Plus Konica Minolta®). The measurement device determines the relative amount of chlorophyll by measuring the absorbance in two wavelength regions where one wavelength corresponds to an absorbance peak of chlorophyll. A numerical SPAD value proportional to the amount of chlorophyll present in the leaf is subsequently calculated from the difference in absorbance. Stomatal conductance was determined with a leaf porometer (SC-1 Leaf Porometer from METER Group ®) which measures the actual vapour flux from the leaf through the stomata out to the environment. Chlorophyll and stomatal conductance measurements were conducted once per site in April, May and June on two subsequent sunny days around midday (SPAD measurements) and during the morning (9-11 am) and afternoon (3-5 pm) hours (porometer measurements) on three individual plants within each plot. Volumetric soil water content measurements were carried out with a ML3 ThetaProbe Soil Moisture Sensor (Delta-T, Cambridge) at two time points (21 April, 26 May). Measurements were taken in the centre of the plots and a standard calibration for mineral soil was used.

At full maturity (BBCH 89), four individual plants in each plot were randomly collected for individual phenotypic trait measurements (plant height, total grain weight, grain mass, number of grains) (22-24 July 2022). The harvested material (ears with short stalks) was threshed with a plot harvester machine “HEGE 150”. Grains were stored in paper bags under dark and dry conditions at room temperature. Thousand kernel weight (TKW) and hectolitre weight were measured with a MARVIN optical grain counter (Digital Seed Analyser, GTA Sensorik GmbH, Neubrandenburg, Germany) and a balance (Mettler PM2000, Mettler-Toledo, Greifensee, Switzerland). Protein (%) and mineral contents (% or mg kg^-1^) were analysed by near-infrared reflectance spectroscopy (NIRS) using a NIRFlex N-500 (Büchi Labortechnik AG). The protein calibration of the NIRFlex was annually adjusted with 50-100 wheat samples from different varieties and origins. Reference analyses were made with the Kjeldahl method, according to ICC standard method No. 105/2. The coefficient of confidentially of the calibration is R^2^ = 0.93 (Cécile Brabant, personal communication). The protein content of the herein examined samples fitted into the range of the NIRS calibration.

### Data Analyses

Statistical analyses were carried out with R version 3.6.1. The data was tested for normality and homogeneity of variance by a visual inspection of residuals (normal quantile-quantile plots, standardized residuals versus fits plot) and revision of coefficients of determination (R^2^). When necessary, logarithmic or square root transformations were conducted to achieve normality and homogeneity of variance. The split-plot design was accounted for by including “longplot” in the random term and a spatial correlation with coordinates of distances from the tree row (“d_cor”). The random term further included a factorial term with site and replicate (“siterep”). The model explaining yield, and plant density, at the plot level was thus: lme(sqrt(yield)∼site*distance*variety, and lme(plantnumber∼site*distance*variety, random=∼1|siterep/longplot, correlation=corGaus(form=∼d_cor)), where “distance” was a factor with four levels and “variety” referred to the variety ‘Wiwa’ and the ‘CC-2k’ population. For better readability, the term “variety” is used in the remainder of this document to refer to both variety and population. At the individual level, “plot” was nested additionally in the random term and a second coordinate (which was randomly assigned to each individual sample) perpendicular to the distance-coordinate had to be given not to create zero distances in the model. Differences in group means among groups was analysed by multifactorial ANOVA (type I, sequential sum of squares). Significances of each factor were assessed by means of the F-test. For post-hoc analysis, the means of treatment groups were compared with a Tukey test (HSD.test()-function within the R agricolae package (de Mendiburu, 2020) with a significance threshold of α = 0.05.

To calculate the relative amount of PAR at different distances from the tree row, the total amount of PAR per day (PPFD in μmol m;^−2^/day) was calculated for each sensor with the formula 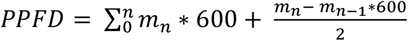, where “m” was the measurement and “n” the number of measurements per day. All measurements were exactly 10 minutes apart (600 seconds) apart. Finally, the sensor placed at 7 m distance to the tree row was taken as the reference and the percental PPFD for the 0 and 3.8 m distance were calculated with the formula 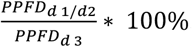. Statistical analyses of chlorophyll content, stomatal conductance and soil moisture followed the model outline above for analyses on the individual level, with the addition of “month” as fixed factor.

## Results

### 1. Plant Density, Yield and Thousand Grain Weight

Plant density was significantly influenced by variety (p < 0.05) and interactions of site and distance (p < 0.001) and site and variety (p < 0.01). In Feusisberg, plant density at 3.8 m from the tree row (41 plants m^-2^) was significantly higher than plant density in Wollerau at 3.8 m (20 plants m^-2^), all other distances had intermediate plant densities (27-40 plants m^-2^). Plant densities of CC-2k and Wiwa were similar in Feusisberg, but statistically different in Wollerau (CC-2k: 45 ± 14 SD plants m^-2^, Wiwa: 16 ± 8 plants m^-2^) (Fig. 2). The correlation coefficient between yield and plant number amounted to 0.70.

**Fig. 2:**
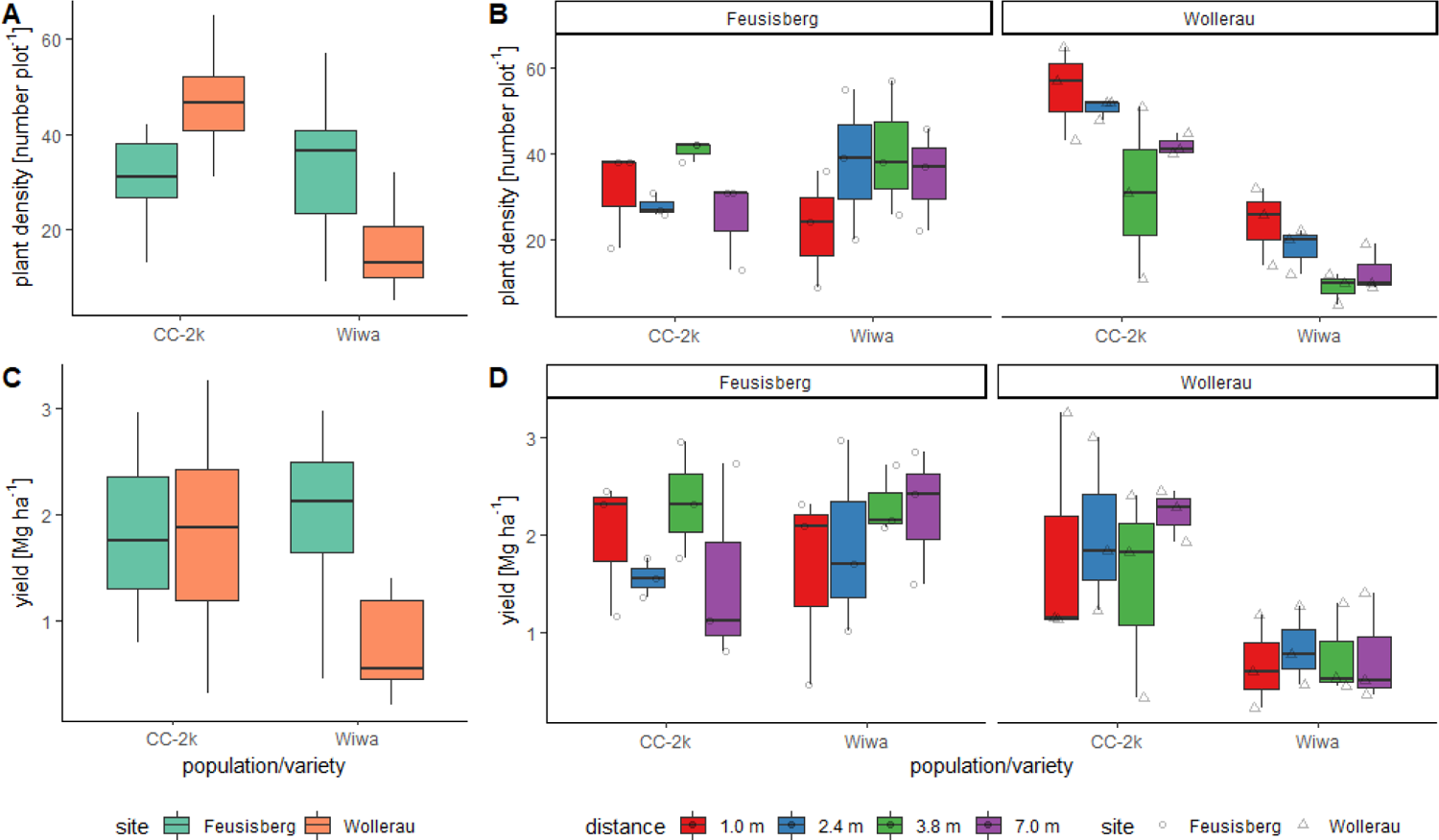
Plant density (A, B) and yield (C, D) of the winter wheat composite cross population ‘CC-2k’ and the common organic variety ‘Wiwa’ at two agroforestry systems in Switzerland. Site-variety effects (A, C) and site-distance effects (B, D) are shown. The box plots range from the first to the third quartile where the horizontal line shows the median. The vertical lines go from each quartile to the minimum or maximum, respectively.

Variety (p < 0.05) and the interactions of site and distance (p < 0.05) and site and variety (p < 0.05) significantly affected wheat yield at the plot level. Yield of CC-2k (1.9 ± 0.8 Mg ha^−1^) was significantly higher than yield of Wiwa (1.4 ± 0.9 Mg ha^−1^) across both sites and all distances. However, CC-2k and Wiwa yield (across all distances) amounted to 1.9 ± 0.7 Mg ha^−1^ and 2.0 ± 0.8 Mg ha^−1^, respectively, in Feusisberg and to 1.9 ± 0.9 Mg ha^−1^ and 0.7 ± 0.4 Mg ha^−1^ in Wollerau, i.e. mean CC-2k yield was identical for Feusisberg and Wollerau while mean Wiwa yield in Wollerau was only 35% of mean Wiwa yield in Feusisberg (Fig). In Feusisberg, yield was highest at 3.8 m (2.3 ± 0.4 Mg ha^−1^) and similar at 1.0, 2.4 and 7.0 m (1.8 ± 0.8, 1.7 ± 0.7 and 1.9 ± 0.9 Mg ha^−1^, respectively). In Wollerau, yield at 1.0, 2.4 and 7.0 m was similar (1.3 ± 1.1, 1.4 ± 0.9 and 1.5 ± 0.9 Mg ha^−1^, respectively), at 3.8 it was lowest (1.1 ± 0.9 Mg ha^−1^). In the post-hoc Tukey test, Wiwa yield in Wollerau was significantly lower than both Wiwa yield in Feusisberg and CC-2k yield in Feusisberg and Wollerau. The interaction of population/variety and distance varied with CC-2k having higher yields in proximity to the tree row in Feusisberg but lower ones in Wollerau compared to the reference yield at 7.0 m distance. For Wiwa, a decreased yield in proximity to the tree row was observed in Feusisberg for the 1.0 and 2.4 m distance, while in Wollerau Wiwa yield was in general very low, amounting to 0.7 to 0.8 Mg ha^−1^ at all distances (Fig. 2). To compare yield variability, the coefficient of variation, also known as relative standard deviation, as the ratio of the standard deviation to the mean was calculated for each variety, being 40% for CC-2k and 64% for Wiwa across the two sites.

Thousand grain weight (TGW) derived from subsamples analysed with MARVIN was significantly affected by distance (p < 0.05). Across both sites, mean TGW was highest at 1.0 and 7.0 m (39.4 ± 3.6 and 39.4 ± 2.9 g, respectively) and lowest at 3.8 m (35.7 ± 5.7 g). The difference between Wiwa (39.0 ± 4.1 g) and CC-2k (37.5 ± 4.1 mg) was not significant. TGW was higher in Feusisberg (39.8 ± 3.1 mg) than Wollerau (36.7 ± 4.5 mg). Variety (p < 0.05) was significant for hectolitre weight with significantly higher weights for Wiwa (77.1 ± 3.4 kg hl^-1^) than CC-2k (73.6 ± 2.6 kg hl^-1^).

The interaction of site and distance was significant (p < 0.0001) for total seed weight per plant and total seed number. Differences in total seed weight per plant were statistically not significant between any distances in Feusisberg but significant between 3.8 m (7.9 ± 4.9 g) and 1.0 m (4.7 ± 4.3 g) in Wollerau. Differences in total seed number per plant were statistically not significant between any distances in Feusisberg but significant between 3.8 m (181 ± 96 seeds) and 1.0 m (98 ± 85) in Wollerau. Distance was marginally significant (p = 0.06) for total seed number with the highest average seed number at 3.8 (151 ± 85), followed by distance 1.0 (131 ± 94), 7.0 (122 ± 63) and 2.4 m (113 ± 58), respectively (across both sites). No effects of site, distance and variety were observed on tillering.

### 2. Quality Parameters and Plant Height

Grain hardness was significantly affected by variety (p < 0.01) and the interaction of variety and distance (p < 0.01) where CC-2k had a higher mean grain hardness (26.3 ± 1.2%) compared to Wiwa (25.0 ± 5.0%). While grain hardness was generally more homogenous in CC-2k and only slightly decreased at 1.0 m distance, it was highest at 1.0 m distance in Wiwa (Fig. 3).

**Fig. 3:**
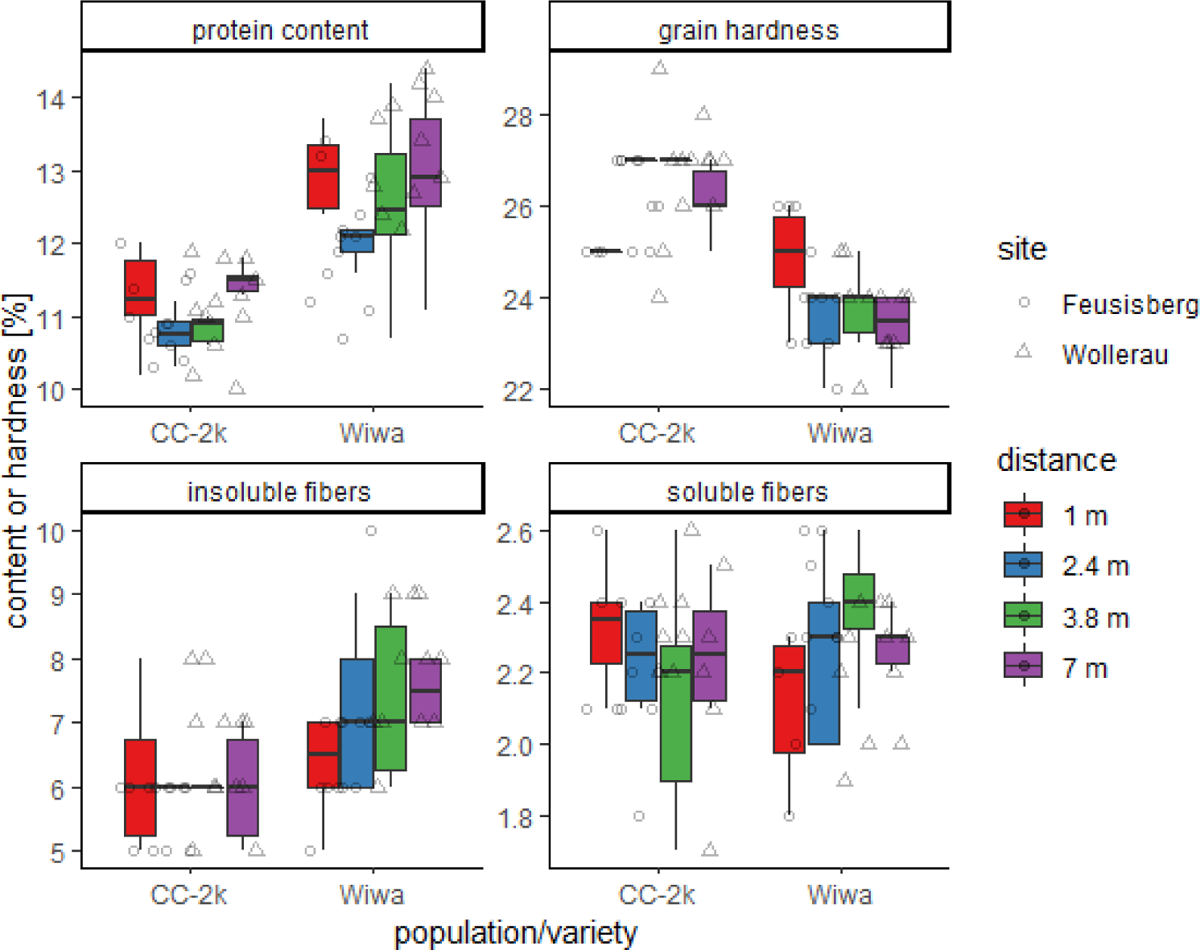
Protein content, grain hardness and content of insoluble and soluble fibers of the winter wheat composite cross population ‘CC-2k’ and the common organic variety ‘Wiwa’ at two agroforestry systems in Switzerland, analysed with near-infrared reflectance spectroscopy (NIRS). The distances refer to the distance to the tree row. The box plots range from the first to the third quartile where the horizontal line shows the median. The vertical lines go from each quartile to the minimum or maximum, respectively.

Protein content was significantly affected by variety (p < 0.01) and marginally significantly by site (p = 0.06). Protein content was significantly higher in Wiwa (12.7 ± 1.0%) than in CC-2k (11.1 ± 0.6%) and in Wollerau (12.2 ± 1.3%) than Feusisberg (11.5 ± 0.9%) (Fig. 3).

Potassium (K), phosphorus (P), calcium (Ca), magnesium (Mg), iron (Fe) and zinc (Zn) content depended on variety (p = 0.07, p < 0.01, p = 0.06, p < 0.01, p = 0.01, p < 0.01, respectively, Fig. 4, see also SI Tab. 10). Some nutrients showed effects on site (Mg), distance (K, P) and interactions of site and variety (Ca, Zn), site and distance (P) and distance and variety (P). Mg content was higher in Wollerau than Feusisberg across both varieties and all distances. For Zn, contents were equal in CC-2k at both sites (Feusisberg: 28.2 ± 1.3 mg kg^-1^, Wollerau: 28.3 ± 1.7 mg kg^-1^) but significantly different between Wiwa in Wollerau (33.6 ± 0.9 mg kg^-1^) and in Feusisberg (30.3 ± 1.9 mg kg^-1^). Wiwa had a significantly higher content of Fe (46.1 ± 4.0 mg kg^-1^) compared to CC-2k (38.5.1 ± 3.3 mg kg^-1^). For Ca, the difference in content between Wiwa and CC-2k was significant in Wollerau (Wiwa: 0.046 ± 0.003%, CC-2k: 0.042 ± 0.003%) but equal in Feusisberg (Wiwa: 0.043 ± 0.002%, CC-2k: 0.043 ± 0.003%). K content was affected by an interaction of site and distance: In Wollerau no significant difference between distances was observed, in Feusisberg at 7.0 m (0.395 ± 0.023%) had significantly higher content than at 1.0 m (0.373 ± 0.010%). Concerning the interaction of distance and variety, no difference between distances were observed for CC-2k while for Wiwa at 1.0 m (0.387 ± 0.015%) had a significantly lower P content than at 3.8 and 7 m (0.407 ± 0.016 and 0.408 ± 0.012%).

**Fig. 4:**
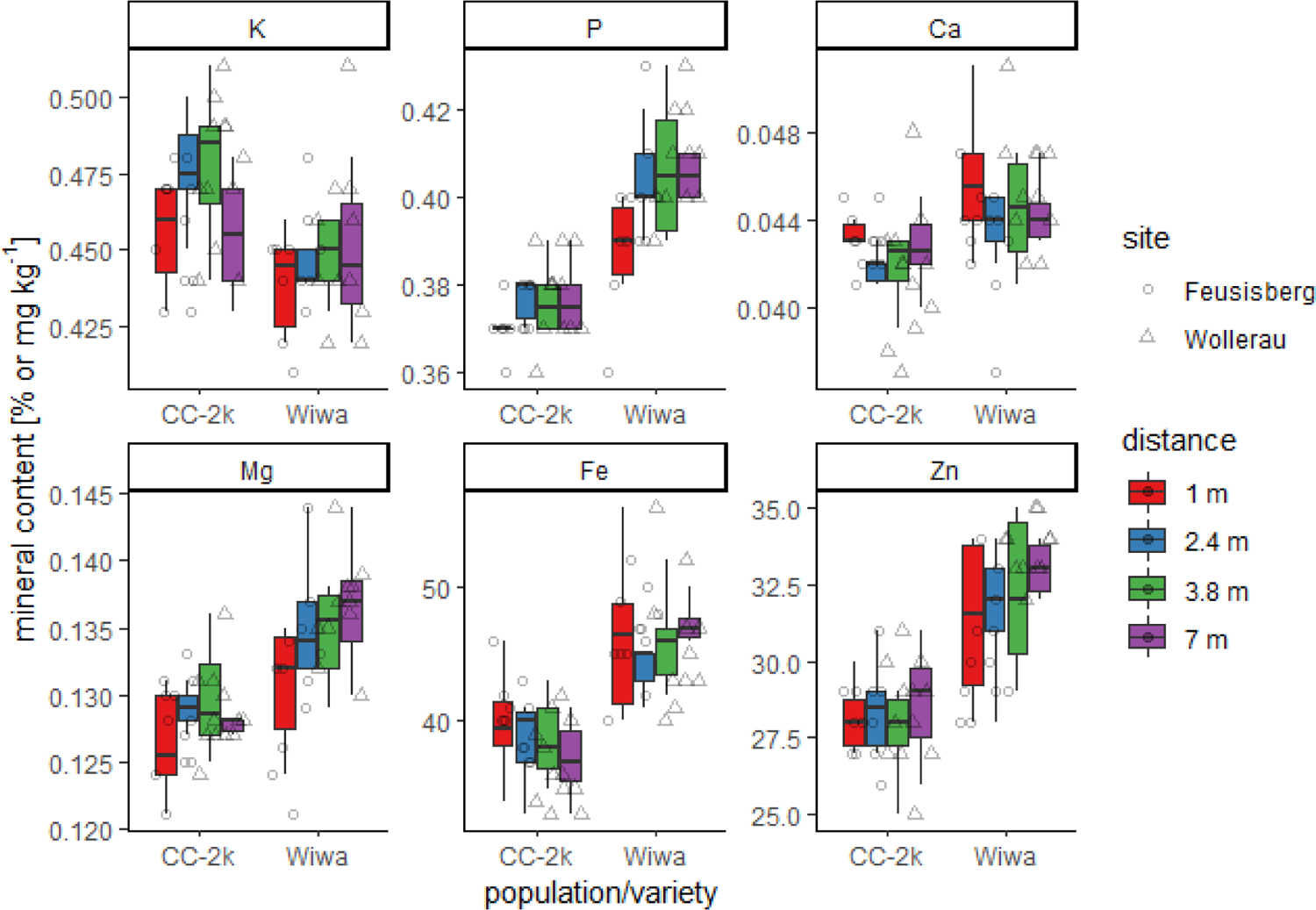
Mineral contents of the winter wheat composite cross population ‘CC-2k’ and the common organic variety ‘Wiwa’ at two agroforestry systems in Switzerland, analysed with near-infrared reflectance spectroscopy (NIRS). Iron (Fe) and Zinc (Zn) contents are given in mg kg^-1^; Potassium (K), Phosphorus (P), Calcium (Ca) and Magnesium (Mg) contents are given in %. The distances refer to the distance to the tree row. The box plots range from the first to the third quartile where the horizontal line shows the median. The vertical lines go from each quartile to the minimum or maximum, respectively.

With respect to soluble and insoluble fibres, CC-2k had the highest soluble fibre content at 1.0 (2.33 ± 0.18%) and the lowest at 3.8 m (2.13 ± 0.033%) while it was the opposite for Wiwa (lowest content at 1.0 m with 2.12 ± 0.21% and highest content at 3.8 m with 2.38 ± 0.17%). This interaction was not significant. For insoluble fibres, site (p < 0.05) and variety (p < 0.05) were significant with higher contents in Wollerau (7.09 ± 1.16%) compared to Feusisberg (6.17 ± 1.05%) and higher in Wiwa (7.17 ± 1.23%) compared to CC-2k (6.08 ± 0.88%).

Distance also influenced plant height significantly (p < 0.001), with plants being higher close to the tree (distance 1-4: 90 ± 10 cm, 88 ± 8 cm, 87 ± 9 cm and 84 ± 10 cm, respectively). Wiwa was marginally significantly (p = 0.05) higher (89 ± 8 cm) than CC-2k (85 ± 10 cm) across both sites. No effects of site, distance and variety were observed on tillering.

### 3. Chlorophyll Content and Stomatal Conductance

Chlorophyll content was significantly affected by month (p < 0.01), site (p < 0.05) and distance (p < 0.01). The interactions of site and distance (p = 0.07) and site and variety (p = 0.07) were marginally significant. Chlorophyll content increased with time with highest values in June (Fig. 5). Feusisberg had significantly higher chlorophyll contents (43.3 ± 4.6) than Wollerau (40.4 ± 4.4). In terms of distance to the tree row, chlorophyll content was higher at 2.4 and 3.8 m (42.9 ± 4.8 and 42.7 ± 4.1, respectively)) than at 1.0 and 7.0 m (41.6 ± 5.2 and 40.3 ± 4.3, respectively) across both sites. In Feusisberg, it was highest at 2.4 m, followed by 1.0, 3.8 and 7.0 m. In Wollerau, it was highest at 3.8 m, followed by distance 2.4, 1.0 and 7.0. In Feusisberg, CC-2k had a higher chlorophyll content (43.7 ± 5.0) than Wiwa (43.0 ± 4.2), though this difference was not significant. In Wollerau, Wiwa had a significantly higher chlorophyll content (41.1 ± 4.0) than CC-2k (39.6 ± 4.6).

**Fig. 5:**
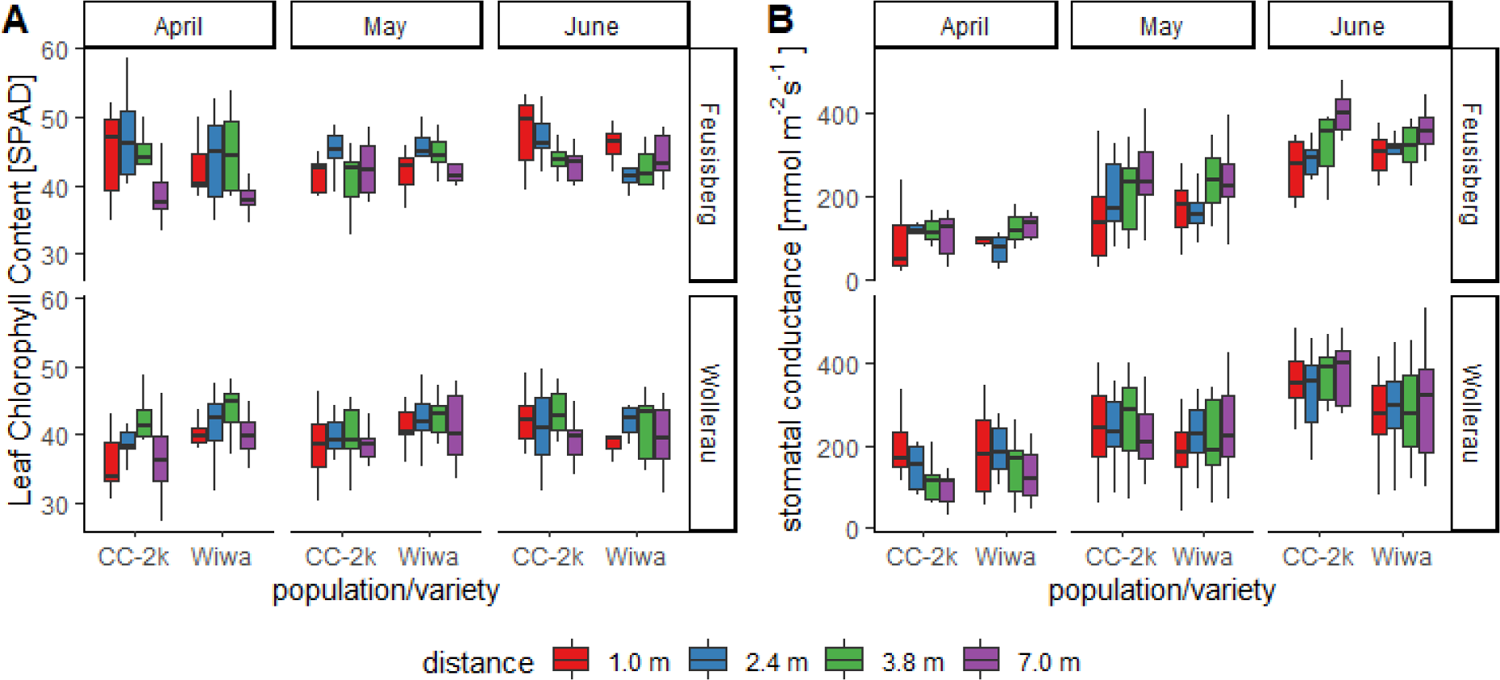
Leaf chlorophyll content (A) and stomatal conductance (B) of the winter wheat composite cross population ‘CC-2k’ and the common organic variety ‘Wiwa’ at two agroforestry systems in Switzerland. Measurements were taken on two subsequent sunny days from April to June. The box plots range from the first to the third quartile where the horizontal line shows the median. The vertical lines go from each quartile to the minimum or maximum, respectively.

Stomatal conductance was significantly determined by month (p < 0.01), distance (p < 0.01) and the interaction of site and distance (p < 0.01). It increased significantly with time (April: 133 ± 70 mmol m^−2^ s^−1^, May: 214 ± 89 mmol m^−2^ s^−1^, June: 314 ± 101 mmol m^−2^ s^−1^). Regarding the site-distance interaction, stomatal conductance was not significantly different between any distances in Wollerau; in Feusisberg stomatal conductance at 7.0 m was significantly higher than at 2.4 and 1.0 m (183 ± 91 and 173 ± 97 mmol m^−2^ s^−1^, respectively) (see SI).

### 4. Photosynthetically Active Photon Flux Density and Volumetric Soil Moisture

The PAR sensors placed at 7 m distance had a higher PPFD than the sensors on the South side of the tree row originally intended as control. Thus, the relative minima, maxima and means of PAR were calculated from division by the sensors placed in 7 m distance to the tree row within each replicate. Across both sites, relative mean was 71 ± 18% PAR at the edge of the tree row and 99 ± 9% PAR in 3.8 m distance, respectively (Fig. 6). At the edge of the tree row, relative maximum was equal to the reference sensor, but relative minimum amounted to 51 ± 23% of it. In 3.8 m distance, relative minimum amounted to 88 ± 18%.

**Fig. 6:**
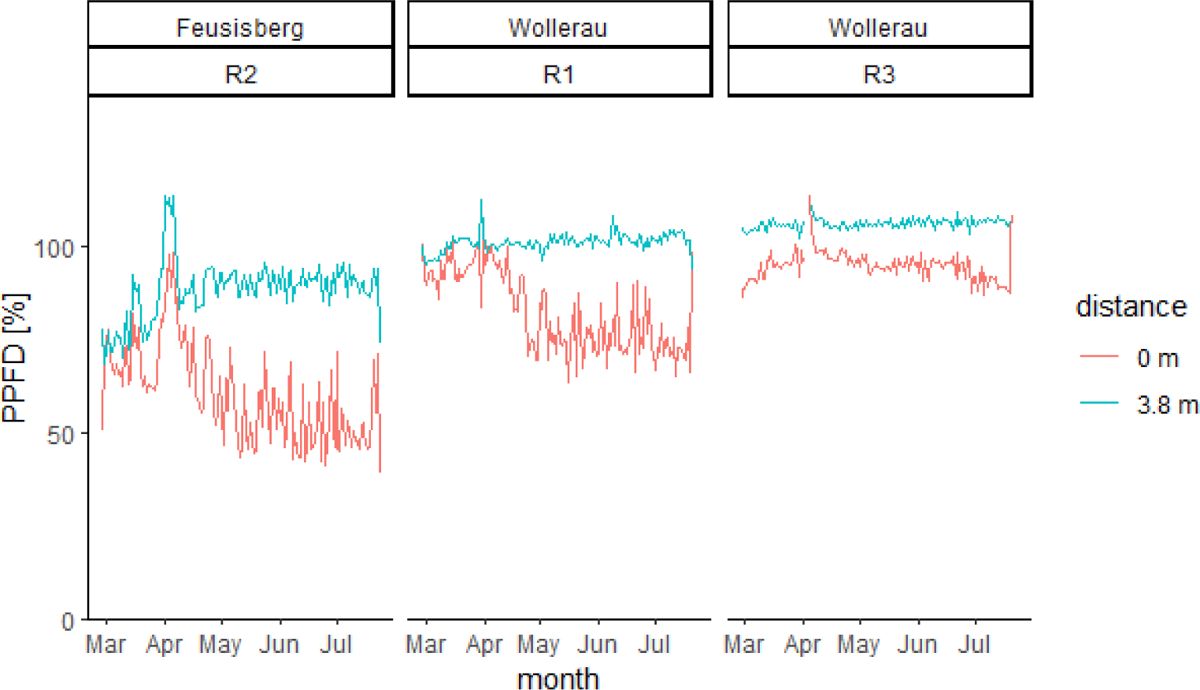
Percentage of Photosynthetically Active Photon Flux Density (PPFD) at the tree row border (0 m distance) and in 3.8 m distance to it at the two experimental sites in Switzerland in comparison to the reference PAR sensor placed at 7 m distance. There was one replicate (R2) with three sensors in Feusisberg and two replicates (R1, R3) with three sensors in Wollerau.

Volumetric soil moisture was significantly affected by month (p < 0.0001) and the interaction of distance and variety (p < 0.05). Distance (p = 0.057) was marginally significant. Soil moisture was significantly higher in May (33.3 ± 5.6%) than April (20.2 ± 8.5%) and at 2.4 m distance to the tree row (31.5 ± 9.6%) than at 1 m (28.3 ± 8.6%) in the post hoc Tukey test. Soil moisture at 3.8 and 7 m distance lay in between. In the CC-2k plots, soil moisture was similar across all distances while the difference between 2.4 m distance (33.9 ± 9.6%) and 1 m distance (27.1 ± 8.8%) was significant.

## Discussion

The main hypothesis of this field experiment was that a genetically diverse cereal population would outperform a high-yielding commercial variety in a heterogenous environment present in an AFS. The underlying assumption is that breeding efforts have created high-yielding modern varieties which are designed for high-input sole crop agriculture, have specific environmental requirements and are not particularly suitable for low-input mixed cropping systems like silvoarable agroforestry (Atlin and Frey, 1989; Smith, Pearce and Wolfe, 2012b; Bourke *et al*., 2021). According to Lammerts van Bueren et al. (2011) 95% of organic production is based on crop varieties bred for conventional agriculture, which has substantially different input levels than organic farming and is relevant for organic breeding (Le Campion et al. 2020). Given that Wiwa is a long established (admission in 2005) and the most commonly grown organic winter wheat variety in Switzerland today (Klaiss and Dierauer 2019), its selection as reference variety was solid. In our field experiment at two agroforestry sites in temperate climate, yield of the CCP CC-2k was more stable across the two sites than yield of the commercial organic variety Wiwa, in the sense that CC-2k showed similar yields in Feusisberg and Wollerau while yield of Wiwa was significantly less in Wollerau than Feusisberg. Also, the coefficient of variation across both sites was lower for CC-2k (40% compared to 64% for Wiwa). More sites are needed, evidently, for a statistically sound coefficient, but it is a first indicator. In terms of yield, CC-2k thus outperformed Wiwa in this trial. A yield-related interaction of variety and distance could not be observed. In comparison to average winter wheat yields from organic production in Switzerland of 4.5 ± 1.3 Mg ha^−1^ in 2022, the achieved yields of 1.9 ± 0.8 Mg ha^−1^ (CC-2k) and 1.4 ± 0.9 Mg ha^−1^ (Wiwa) across all distances and both sites are low (Agristat, 2022). Even the higher yields at 7 m distance to the tree row (2.2 ± 0.3 Mg ha^−1^ for CC-2k and 2.3 ± 0.7 Mg ha^−1^ for Wiwa) were only half of average yield in Switzerland. We attribute various reasons for this circumstance: relatively high losses due to the use of old machinery equipment (seeder and thresher) with small plot sizes, weed pressure and bird predation. This may have mitigated differences between plot distances near the tree row and the reference plot at seven metres which would have been expected (as they occurred, e.g., in Pardon *et al*., 2018). At the same time, tree height was maximum 6 m and trees were positioned 1.3-2.3 m from the border of the tree row. Plots at 7 m distance to the tree row could have been shaded only for a very short period of time at dawn and dusk. PAR sensor data further showed that even 4 m from the tree row light conditions were very similar to the reference sensor at the 7 m distance position. The results should therefore not be interpreted to mean that shading in AFS was the cause of the lower yields. Indeed, the small magnitude of the differences between the distances in all parameters suggests that other factors were of greater importance than the influence of the tree row. Plant density is of major importance to maximise yield (Bastos *et al*., 2020) and final plant density in our study was clearly below the initial seeding density of approx. 350 seeds per m^2^. Though in high-yielding environments, the agronomic optimum planting density is considerably lower than in low-or medium-yielding environments, still its estimates range from 140 to 400 plants per m ^2^ (Bastos *et al*., 2020), being well above the attained plant densities in our trial.

Most wheat parameters showed no effects regarding the distance to the tree row. In contrast, effects (i.e., reductions) of light and soil moisture were observed for the 1 m distance. However, shading and competition for water were not really reflected in the wheat. There was also no difference in wheat parameters at the 2.4 m distance, where soil moisture was apparently increased by shading, or at the area outside the influence of the tree. The fact that both soil moisture and light changed with distance, but the wheat parameters hardly changed at all, is interesting and can be interpreted as a certain physiological plasticity to cope with heterogeneous conditions in wheat. As shown in a 40-year long experimental trial, varieties which performed well at high-input levels were also the best at low input levels, suggesting that interactions between the genotype and environmental conditions were not strong enough to inverse the performance ranking (Büchi *et al*., 2016). Overall, CC-2k and Wiwa demonstrated similar plasticity in terms of distance-effects in this trial.

TGW commonly decreases with shade (e.g., Qiao *et al*., 2019; Artru *et al*., 2017; Li *et al*., 2010). Other studies have reported increased (Zhang *et al*., 2022) or similar (Vaccaro, Six and Schöb, 2022; Zhang *et al*., 2022) TGWs under moderate shade. In this study, TGW was below average at both sites (gzpk 2022, Agroscope, pers. communication), and highest at 1 and 2.4 m distance for CC-2k and Wiwa, respectively, and lowest, for both, at 3.8 m distance. However, PAR measurements showed that the mean reduction in PPFD at 3.8 m distance was minor. Apart from light, temperature, water availability and micronutrients may affect TGW (Arif *et al*., 2006; Kaur and Behl, 2010). Soil moisture measurements did not indicate competition for water at 3.8 m distance, though, and neither P nor micronutrient contents showed reduced values in 3.8 m distance. Total seed weight per plant was also lowest at 3.8 m, while total seed number was highest there. The negative relationship between seed number and weight is well-known (Knott and Talukdar, 1971) and was similar for CC-2k and Wiwa.

Protein content as important parameter for baking quality was slightly lower in CC-2k, though still in the range of minimum protein content for white flour which was typically specified for 10-12% (Kamel and Stauffer 1993). Nowadays it is known that the relationship between protein content and bread volume drops above 12% protein content (Gabriel *et al*., 2017). In a study over a 5-year-period, baking quality of 11 winter wheat varieties were assessed and the influence of year was on a higher level for all evaluated baking quality characteristics compared to the influence of variety – with the influence of year being most important to protein content (Koppel and Ingver 2010). Some quality parameters were slightly higher in Wiwa (protein content, Fe, insoluble fibres) but also showed interactions with site (e.g., Ca where the difference between Wiwa and CC-2k was significant in Wollerau but equal in Feusisberg). Most mineral nutrient contents were higher in Wiwa, though. K content was affected by an interaction of site and distance but not variety. P content was affected by an interaction of variety and distance with no observable differences between distances for CC-2k – supporting for this variable the hypothesis that CCPs can buffer tree impacts. However, P content is of minor relevance for grain quality.

The significant interaction of site and variety, which was of greater importance than distance-effects in terms of yield and mineral nutrient content, may be explained by the lower soil fertility in Wollerau. Besides the lower C:N ratio (8.8) observed in soil samples, leaf measurements with the chlorophyll meter suggest significantly lower nutrient contents. The CC-2k as a CCP proved to be more suited to cope with lower nutrient availability in this trial, as proposed by plant breeders (Atlin and Frey, 1989; Döring *et al*., 2011, 2015; Smith, Pearce and Wolfe, 2012a; Bourke *et al*., 2021).

Lastly, there is the aspect of natural adaptation over time of CCP grown year after year in the same location and the evolution of facilitative interactions, which should lead to higher and more stable yields (Schöb, Brooker and Zuppinger-Dingley, 2018; Vollenweider and Spieß, 2018). Improvements can certainly be expected compared to varieties bred in highly fertile, weed-free, densely seeded environments (Atlin and Frey, 1989). Besides that, the importance of maintaining and enhancing agrobiodiversity and diverse genetic varieties is emphasized and encouraged by major agreements and other policy instruments such as the FAO’s International Treaty on Plant Genetic Resources for Food and Agriculture and the Nagoya Protocol on Access to Genetic Resources and the Fair and Equitable Sharing of Benefits Arising from their Utilization.

## Conclusion

In summary, the CC-2k was more apt to grow well under the heterogeneous and low-input conditions at both agroforestry sites and outperformed the commercial organic variety Wiwa in terms of yield but not in terms of grain quality. As variety-/population-specific yield was independent from the distance to the tree row, it may be concluded that shade-induced reductions in yield-related characteristics must not have been decisive or might have been outweighed by positive interactions or a beneficial microclimate in the AFS of this experiment.

## Declaration

This work was supported by the World Food System Center [Call 2 | 2021].

## Acknowledgments

We thank Karl Eggler and Jakob Bürgi von Aarburg for their permission and help to conduct a field experiment on their fields. We are particularly grateful for Stef den Hond and his continuous physical and mental support throughout the experiment. We also thank Anita Vaccaro, Helga Palli, Maria Schubert-Kastner, Johannes Blacha, Levin Spiegel and Clemens Proksch for their help during after harvest.

